# Tumorigencity decrease in Bcl-xL deficient MDCK cells ensuring the safety for influenza vaccine production

**DOI:** 10.1101/2024.09.14.613056

**Authors:** Jiahao Zheng, Boran Li, Lanxin Jia, Jiayou Zhang, Zheng Gong, Yang Le, Xuanxuan Nian, Xuedan Li, Bo Liu, Daiguan Yu, Zhegang Zhang, Changgui Li

**Affiliations:** National Engineering Technology Research Center for Combined Vaccines, 430207, Wuhan, China; Wuhan Institute of Biological Products Co.Ltd., 430207, Wuhan, China; Medical Products Administration of Hubei Province, 430071, Wuhan, China; Division of Respiratory Virus Vaccines, National Institute for Food and Drug Control, Beijing, China

**Keywords:** Madin-Darby Canine Kidney cell (MDCK), Bcl-2-like protein 1 (Bcl-xL), Apoptosis, Tumorigenicity, CRISPR-Cas9

## Abstract

Madin-Darby canine kidney (MDCK) cells are the recognized cell strain for influenza vaccine production. However, the tumorigenic potential of MDCK cells raises concerns about their use in biological product manufacturing. To reduce MDCK cells’ tumorigenicity and ensure the safety of influenza vaccine production, a B-cell lymphoma extra-large (Bcl-xL) gene, which plays a pivotal role in apoptosis regulation, was knocked-out in original MDCK cells by CRISPR-Cas9 gene editing technology, so that a homozygous MDCK-Bcl-xL^-/-^ cell strain was acquired and named as BY-02. Compared with original MDCK cells, the proliferation and migration ability of BY-02 were significantly reduced, while apoptosis level was significantly increased, the endogenous mitochondrial apoptotic pathway were also modulated after Bcl-xL knock-out in MDCK cells. For tumor formation assays in nude mouse tests, all ten mice injected with original MDCK cells presented tumors growth in the injection site, in contrast to only one mouse injected with BY-02 cells presented tumors growth. These findings suggest that Bcl-xL knock-down is an effective strategy to inhibit tumor formation in MDCK cells, making BY-02 a promising genetically engineered cell strain for influenza vaccine production.

## 1. Introduction

Globally, influenza virus causes an estimated 3–5 million severe cases and 300 000–650 000 deaths each year, which places a heavy economic burden on low- to middle-income countries [1,2]. The influenza virus is a single-stranded, segmented RNA virus with low RNA polymerase fidelity, and thus, its genomic mutation rate significantly increases relative to other viruses [3]. Seasonal influenza vaccine is crucial to deal with the global influenza epidemic. The eggs laid by embryonated hens have been used as substrates to produce influenza vaccines for >60 years. Influenza virus often acquires antigenic changes through host adaptation during serial passages in eggs. The most common amino acid sequence changes are observed in hemagglutinin and neuraminidase, which reduce the protective effect of the vaccine [4–6]. Studies have shown that egg adaptation can lead to a 7%–21% reduction in vaccine match (VM) and a 4%–16% reduction in influenza vaccine effectiveness (IVE). This is similar to the effect of antigen drift on VM (8%–24%) and IVE (5%– 20%) [7]. Since its establishment in 1958, the Madin-Darby canine kidney (MDCK) cell line has been extensively used for amplifying and purifying various viruses. Due to their high susceptibility, rapid proliferation and stable passage matrix for the influenza virus, MDCK cells are recognized as one of the most suitable cells for the production of the influenza virus vaccine [8,9]. However, Intravenous inoculation of chicken embryos with MDCK cells has been previously induced tumor occurrence in the embryonic brain and chorioallantoic membrane nodules [10]. The nodules were observed in neonate BALB/c nude mice inoculated with MDCK cells [11], and they were found to be adenocarcinomas using histological analysis [12]. In other studies, MDCK cells adapted for suspension culture caused tumor formation only injecting with as few as ten cells per nude mouse [13]. Thus, to improve the safety of the influenza vaccine, the best solution is to develop nontumorigenic MDCK cell strains.

Apoptosis is an important part of programmed cell death and is the main mode of regulating the steady state of cell populations. Abnormal apoptosis can lead to neurological diseases [14], immune system abnormalities [15], and cancer [16]. Cancer is a classic example of abnormal cell cycle regulation where excessive cell proliferation or decreased cell removal occurs [17]. The inhibition of cell apoptosis is the main mechanism of certain cancers. Apoptosis mainly includes the intrinsic apoptotic pathway or mitochondrial pathway and the extrinsic apoptotic pathway or death receptor pathway [18]. The regulation and control of the mitochondrial apoptosis pathways are mainly implemented by the Bcl-2 family members [19], and the imbalance of apoptosis with anti-apoptosis proteins causes several cancers. Tumor cells can promote anti-apoptotic protein expression or downregulate proapoptotic protein expression to avoid programmed death. Bcl-xL, an anti-apoptotic protein, has been recognized as an important factor regulating the mitochondrial pathway, and it shows high expression in numerous cancer cells [20]. Inhibiting Bcl-xL efficiently triggers apoptosis of melanoma [21] and ovarian cancer cells [22].

The mechanism of the effect of Bcl-xL on the tumorigenicity of MDCK cells remains unclear. Therefore, in this study, we used CRISPR-Cas9 technology to construct a Bcl-xL deletion cell strain named as BY-02 and analyzed the effect of Bcl-xL on MDCK cell proliferation, migration, and tumorigenicity. Its molecular mechanism was explored using transcriptomics. The aim of the study was to identify a safe and reliable genetic engineering cell strain for influenza vaccine production.

## 2. Materials and Methods

### 2.1 Cell lines and cultivation conditions

The original MDCK (CRL-2935TM) cells were bought from the American Type Culture Collection (ATCC, USA). MDCK cells were cultured in VP-SFM (Gibco) medium containing 5% FBS (Gibco) and 1% glutamine. 2BS, VERO cells (ATCC, USA) were cultured in DMEM (Gibco) containing 5% FBS (Gibco) and 1% glutamine, followed by inoculation in T-flasks (Corning) and incubation at 37°C under 5% CO2.

### 2.2 Mice

The 4–7-week-old female nude mice were provided by Vital River Laboratory Animal Technology Co., Ltd. (Beijing) for cell tumorigenicity analysis. All the animal experimental protocols received approval from the Ethical Review Committee for Experimental Animal Welfare of Wuhan Institute of Biological Products Co., Ltd. (protocol code WIBP-AII312023002, 22 July. 2022). Our study with observational experimental design was carried out in compliance with the ARRIVE guidelines.

Anesthesia method: Inhalation anesthesia. Anesthetic name: Isoflurane. Method of execution: Cervical dislocation and death.

### 2.3 Construction of bcl-xl knockout with CRISPR-Cas9 technology and cell screening

The *bcl-xl* gene of the corresponding cannine species was found in NCBI, and the single-guide (sg) RNA sequence with high specificity located in exon 1 was screened through the CRISPR design website (http://www.e-crisp.org/E-CRISP/). The gRNA sequence was synthesized and cloned in pX459 plasmid harboring Cas9. Meanwhile, a targetting vector containing the upstream and downstream homology arms at the target site and the puromycin selection marker was constructed. Half million of MDCK cells in a cuvette were electroporated with 8 µg plasmids containg the pX459 and the targetting vector with the Gene Pulser Xcell electroporation system (Bio-Rad, USA) at 130 V, 1 pulse, and 25 ms pulses . The cells were then incubated at 37°C under 5% CO_2_. The medium was changed at 48-h post-electroporation and then the selection of cell clones was selected with the complete medium containing 8 µg/mL puromycin. Monoclonal cells were obtained by limiting dilution assay.

### 2.4 Real-time quantitative reverse transcription PCR (qRT-PCR)

Total RNA from MDCK cells was extracted with Trizol reagent and quantified with the microplate reader (Thermo Fisher, USA). The total RNA was used as a template for reverse transcription to cDNA according to the specification of PrimeScriptTM IV 1^st^ strand cDNA Synthesis Mix (TaKaRa, Japan). Next, the amplification of DNA was according to the specification of ChamQ SYBR qPCR Master Mix (Vazyme, China) by 7500 Fast Real-Time PCR System (ABI Life Technologies, Singapore). was employed for recording. The quantify of the target gene was determined by compared to the β-actin internal reference.

### 2.5 Western-blotting

After washing with prechilled PBS, the MDCK cells were lysed with the buffer containing 50 mM Tris-HCL, 1% TritonX-100, 150 mM NaCl, 1% sodium deoxycholate, 0.1% SDS, and 100x protease inhibitor. Protein quantification was performed using the Lowry method. Protein samples were separated via electrophoresis using 8% SurePAGETM (GenScript) gels, transferred to nitrocellulose membranes and subjected to immunoblotting. The membranes were then blocked overnight using 5% bovine serum albumin at 4°C, followed by 2 h incubation with primary antibody (1:1000) under ambient temperature. Membranes were washed with Tris-buffered saline with Tween 20 and incubated with HRP-labeled goat anti-rabbit or anti-mouse IgG secondary antibody (Sangon; 1:10000) for 2 h. Amersham ImageQuant 800 system (Cytiva, Japan) was utilized for scanning and analysis. The proteins on the membrane were detected using the following specific antibodies: anti-Bcl-xL (Proteintech Cat. 10783-1-AP), anti-Bax (Sangon Biotech Cat. D197138), anti-Cyto-C (Servicebio Cat. GB11080), anti-Caspase 3 (Servicebio Cat. GB11767C), anti-VDAC1 (Sangon Biotech Cat. D124100) and anti-ACTB (Sangon Biotech Cat. D110001).

### 2.6 Cell proliferation and metabolic levels

Following trypsin digestion, MDCK cells in the logarithmic phase were collected and then washed thrice with PBS. Approximately, 2.5 × 10^5^ cells were seeded into T25 flasks and cultured at 37°C under 5% CO_2_. Thereafter, the medium composition and the cell count were recorded at 0, 12, 24, 36, 48, 60, 72, 84, 96, 108, and 120 h following inoculation. The cell growth and metabolic curves were drawn with the average value of three repetitions for each group of cells.

### 2.7 Cell migration assay

MDCK cells in the logarithmic phase were collected and washed as described above. 2 × 10^4^ cells were seeded into Incucyte Woundmaker 96-well Rinse Boat Assemblies (Sartorius) and cultured with Incucyte® SX5 Live-Cell Analysis System at 37°C under 5% CO_2_. Once the cells reached 100% confluence, Incucyte Woundmaker Tool (Cat.No. 4563) was used to make a scratch. After washing with PBS, images were collected at 2h intervals and measured with Image J to calculate the width.

### 2.8 Apoptosis assay

MDCK cells in the logarithmic phase were collected and washed as described above. Approximately, 1 × 10^6^ cells were resuspended in the binding buffer, followed by a 15 min incubation with Annexin V-APC and propidium iodide (PI) in the dark at ambient temperature using the Annexin V-APC/PI Apoptosis assay kit (Procell) and detected using flow cytometry.

### 2.9 Mitochondrial isolation

The collected MDCK cells were rinsed twice with prechilled PBS. Next, 1 mL of cytoplasm extraction buffer (1 µL protease inhibitor, 5 µL phosphatase inhibitor, and 1 µL DTT added before use) was added. The cells were homogenized 50 times on ice using a homogenizer to obtain the cell fragmentation rate more than 90% exanimated under microscopy. Mitochondria were precipitated via centrifugation at 12000 rpm for 30 min at 4°C. Subsequently, the precipitate was resuspended with 100 µL of cytoplasm extraction buffer for 30s and centrifuged again at 12000 rpm for 10 min at 4°C, and the supernatant was discarded. Mitochondrial lysis buffer was added to the precipitate and left on ice for 30 min, followed by centrifugation at 12000 rpm for 10 min at 4°C the mitochondrial supernatant was collected.

### 2.10 Tumor xenograft studies

The 4-7-week-old female nude mice were classified into four groups: the positive control group injected with 1 × 10^6^ HeLa cells per mouse; the negative control group injected with 1 × 10^6^ 2BS cells per mouse; the control group injected with 1 × 10^7^ WT MDCK cells per mouse; the experimental group injected with 1 × 10^7^ BY-02 cells per mouse. The cells with ≥90% viability were suspended in prechilled PBS and followed with subcutaneous injection on the back of the mouse. Mouse body weight was monitored during the experiment, and the tumor volume was monitored using caliper measurements every three days. Mice were sacrificed after the tumor size reached an ethically unacceptable volume, followed by tumor tissue dissection for subsequent analyses.

### 2.11 RNA sequencing of the transcriptional profile of MDCK cells

Total RNA was extracted from WT cells and BY-02 cells and sent to Sangon Biotech (Shanghai) for RNA sequencing. The main processes included: raw data quality control, transcriptome assembly, gene annotation, expression level analysis, expression difference analysis, and gene enrichment analysis (Supplementary file 3).

### 2.12 Statistical analysis

Data were represented as mean ±SEM and analyzed using GraphPad Prism 9.0. An unpaired t-test was used for comparisons between the two groups. * p<0.05, ** p<0.01, and *** p<0.001 represented statistically significant differences.

## 3. Result

### 3.1 Construction of MDCK cell strain with Bcl-xL deficiency

The already-validated CRISPR system to make homozygous deletion in MDCK cells was employed [23]. To achieve complete deficiency, we formulated the “targeting the key domain structure plus frame shift mutations” strategy. Bcl-xL contains four BH (Bcl-2 homology) domains which have a critical effect on the tertiary structure. Besides, the BH1–BH3 domains form the hydrophobic pocket that interacts with other BH3 domain in proapoptotic proteins to exert an anti-apoptotic function [24]. A gRNA sequence was selected to target the BH3 domain sequence located closer to the start codon on exon-1, which could create the premature appearance of a termination codon with a frame-shift mutation (Fig. 1A). It was obtained a MDCK cell clone with Bcl-xL homozygous deletion, MDCK-Bcl-xL^-/-^ named as BY-02. BY-02 cells were determined with Bcl-xL transcription and protein expression levels. RT-PCR analysis (Fig. 1B) showed a marked decrease in the Bcl-xL mRNA expression levels in BY-02 cells, and western blot analysis (Fig. 1C) showed no Bcl-xL specific band. This indicated that Bcl-xL had been completely and successfully knocked out.

**Fig. 1.**
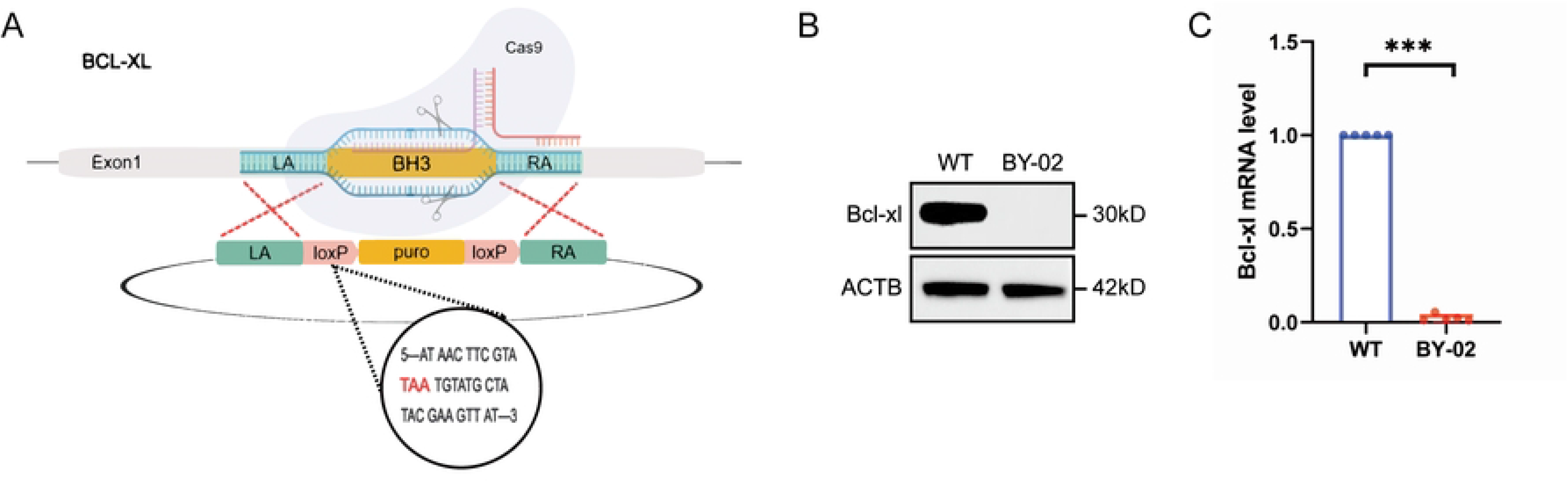
The targeting strategies, qRT-PCR, and western bolt to verify BY-02 cells. (A) The BH3 sequence of *bcl-xl* was replaced by the puromycin resistance gene sequence (puro). LA/RA represents upstream and downstream homologous sequences, and yellow marks the substituted BH3 sequence and puro sequence, respectively. The loxP fragment contains stop codons that prematurely terminate protein translation. (B) Protein level of Bcl-xL in BY-02 cells was determined by Western blotting, and ACTB was used as an internal control protein. (C) qRT-PCR analysis on *bcl-xl* mRNA expression in WT and BY-02 cells. The experimental data are expressed as "mean ± standard deviation". *P < 0.05 and **P < 0.01 indicate significant differences, and ***P < 0.001 indicates extremely significant differences. Data are representative of at least three independent experiments.

### 3.2 Bcl-xL deficiency inhibited MDCK cell proliferation and migration in vitro

To ascertain the impact of Bcl-xL on MDCK cell proliferation and migration, the growth curve of BY-02 cells was first plotted using cell counts (Fig. 2A). A decrease in the proliferative activity of BY-02 cells was observed compared to WT cells, extending the doubling time from 17.56 to 23.43 hours. The cell culture medium’s supernatant was analyzed at 12-hour intervals (Fig. 2B–H), revealing that both BY- 02 and control cells rapidly metabolized glucose, producing lactate and ammonium ions without significantly altering salt ion (potassium, sodium, and calcium) concentrations. Further experiments, including plate clonogenesis and scratch tests (Fig. 2I and J), were conducted. These tests demonstrated that the BY-02 cells had a considerable reduction in the clonogenic ability (Fig. 2K) and a decrease in cell migration rate by approximately 10% compared to WT cells (Fig. 2L). These findings showed that Bcl-xL deficiency markedly inhibited the proliferation and migration of MDCK cells in vitro, indicating Bcl-xL could play a role in reducing tumor formation capacity in MDCK cells in vivo.

**Fig. 2.**
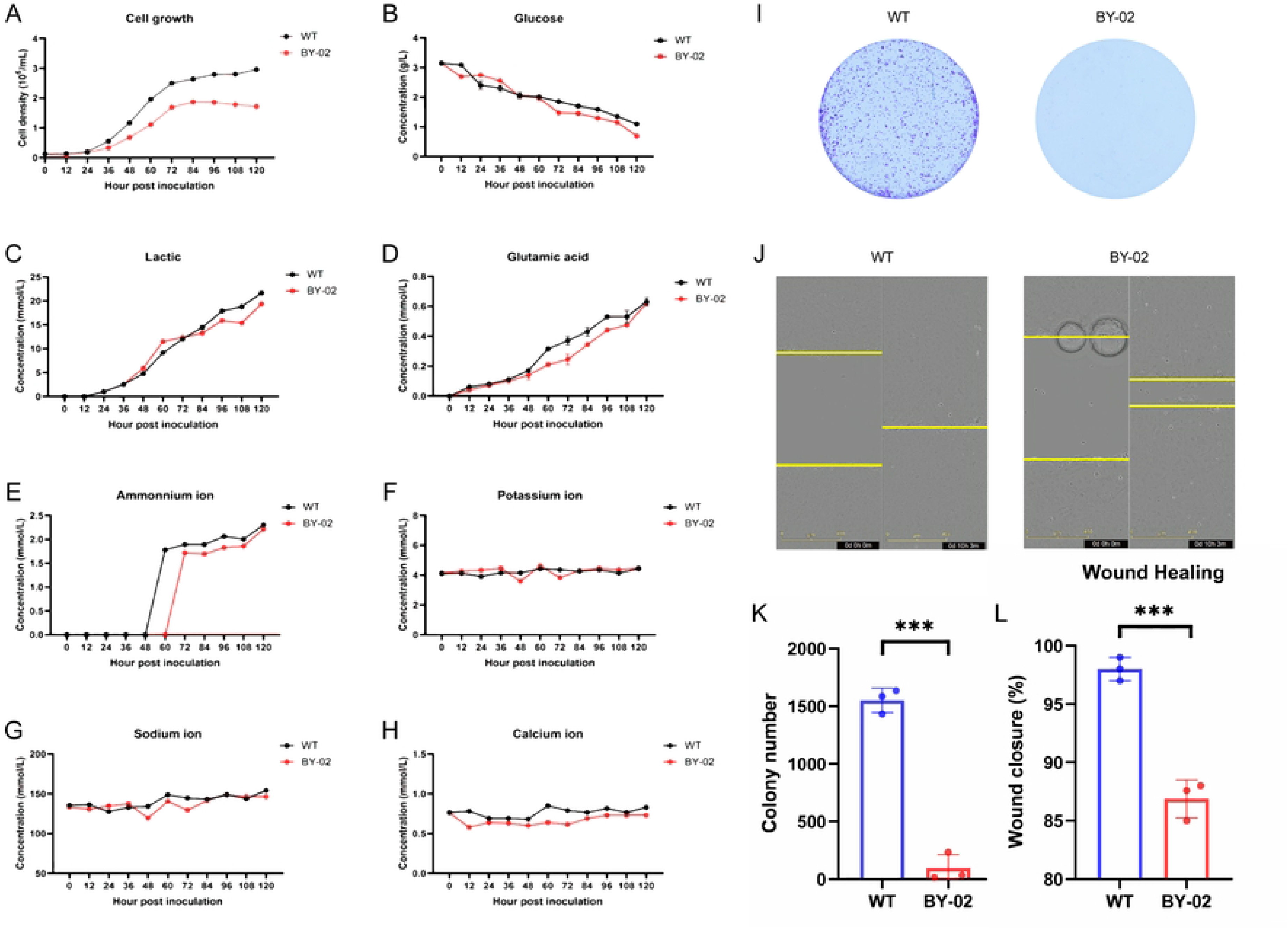
Differences in growth, metabolism and migration of BY-02 cells compared with WT cells. (A) WT and BY-02 cells were counted at 12h intervals to detect differences in cell proliferation. (B-H) The metabolic capacity of WT and BY-02 cells was measured at 12h intervals. (I,K) The effect of Bcl-xL depletion on colony formation rate of MDCK cells. (J,L) The effect of Bcl-xL deletion on MDCK cell migration was detected by scratch assay. The experimental data are expressed as "mean ± standard deviation". *P < 0.05 and **P < 0.01 indicate significant differences, and ***P < 0.001 indicates extremely significant differences. Data are representative of at least three independent experiments.

### 3.3 Inhibition of tumorigenicity in BY-02 cells

The previous in vitro experiments confirmed that Bcl-xL deletion affected the proliferation and migration in MDCK cells. The tumorigenic ability of WT and BY- 02 cells was next evaluated in vivo through xenografts performed in nude mice. 1 × 10^7^ cells were inoculated into the posterior neck of nude mice, and the tumors were dissected four months after inoculation (Fig. 3A). Subcutaneous tumors were observed in all ten mice which received WT cell injections. In contrast, only one of ten mice which received BY-02 cell injections developed subcutaneous tumor (Fig. 3B) even with a significant reduction in tumor size compared to the WT group (Fig. 3C). All mice underwent dissection, and the major tissues (heart, liver, spleen, lung, and kidney) as well as tumors were sectioned and stained with hematoxylin & eosin for pathological analysis (Fig. 3D).

**Fig. 3.**
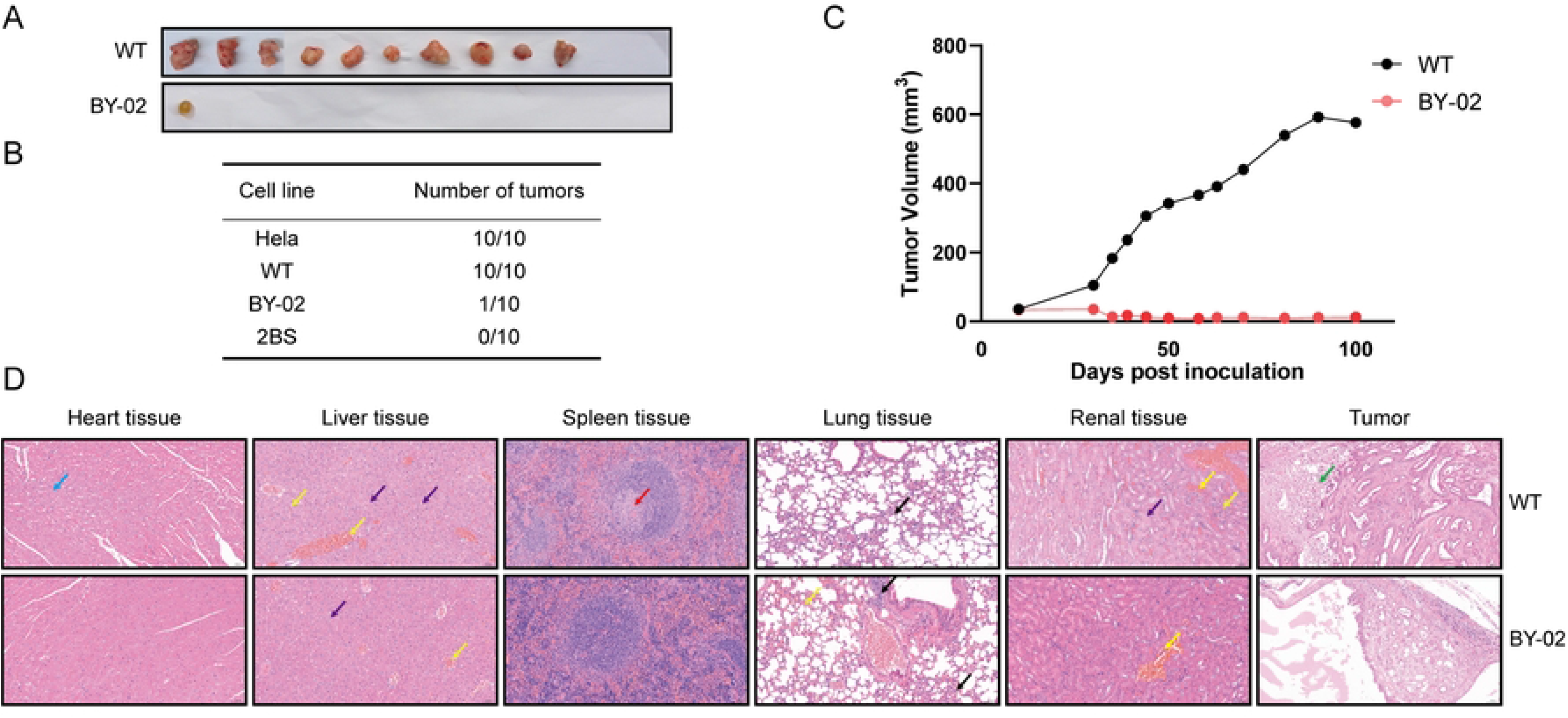
Effect of Bcl-xl on the tumorigenicity of MDCK cells in nude mice. (A) After subcutaneous injection of WT and BY-02 cells into the posterior neck of nude mice, nude mice formed tumors. BY-02 cells showed lower tumorigenicity than WT cells. (B) Statistical results of subcutaneous tumor number in nude mice. In 10 mice, WT cells formed 100% tumors, while BY-02 cells only formed tumors in one mouse. (C) Tumor volume comparison. The tumor volume formed by BY-02 cells in nude mice was much smaller than that of WT cells. (D) Histological observation of heart, liver, spleen, lung, kidney and tumor of nude mice 100 days after subcutaneous injection of WT and BY-02 cells (H&E staining). Myocardial cells showed vacuoles in the cytoplasm (blue arrows), hydropic degeneration of the cells, swelling of the cells, loose and pale staining of the cytoplasm (purple arrows), infiltration of lymphocytes and neutrophils (black arrows), and focal necrosis (green arrows) with a small amount of congestion (yellow arrows) in the tissue.

The control group after injection of WT cells, Cardiac tissue exhibited an increased number of cytoplasmic vacuoles. Most hepatocyte samples displayed degeneration and necrosis, accompanied by inflammatory cell infiltration. Most of the samples showed few cells surrounding the central artery of the spleen’s white pulp. Lymphocytes and neutrophils infiltrated lung tissue, and the alveolar wall displayed slight thickening. There was an increased number of samples with renal tissue congestion, and a small number of samples showed hydropic degeneration of renal tubular epithelial cells.Tumor tissue exhibited large columnar or cuboidal cells arranged in a glandular tubular pattern, with extensive areas of necrosis observed in most samples. In contrast, mice injected with BY-02 cells showed no severe symptoms and were in good condition. These findings indicated that the knockout of Bcl-xL inhibited the formation of tumors in nude mice by MDCK cells and MDCK cell tumor metastasis did not occur within 100 days following injection.

### 3.4 Bcl-xL deficiency promoted MDCK cell apoptosis through the mitochondrial pathway

Bcl-xL belongs to the Bcl-2 protein family and plays an anti-apoptotic role. Its deficiency leads to increased apoptosis levels [25]. Annexin V and PI double staining was used to detect the degree of apoptosis in this test. It worked as following when apoptosis occurs, phosphatidylserine on the internal cell membrane binds to Annexin V and PI will stain the late apoptotic and necrotic cells. It showed the apoptosis rate of BY-02 cells reached 40% (compared to WT cells) as analyzed by flow cytometry (Fig. 4A). The proteins involved in the mitochondrial apoptotic pathway (Fig. 4B) were examined, which showed that the internal balance of cells was disrupted following Bcl-xL loss. Western-blotting analysis revealed the expression level of proapoptotic protein Bax was upregulated and a significant increase in the amount of cytochrome C in BY-02 cells. ELISA showed an approximately two-fold increase in the amount of cytochrome C secreted into the supernatant (Fig. 4C). Next, mitochondria were extracted from the cells to detect changes in Bax level in the mitochondria and cytoplasm. The results showed that the deletion of Bcl-xL increased the localization of Bax in the mitochondria in BY-02 cells (Fig. 4D). Altogether, the loss of Bcl-xL initiated the mitochondrial apoptotic pathway and was critical for the increased apoptotic rate in BY-02 cells.

**Fig. 4.**
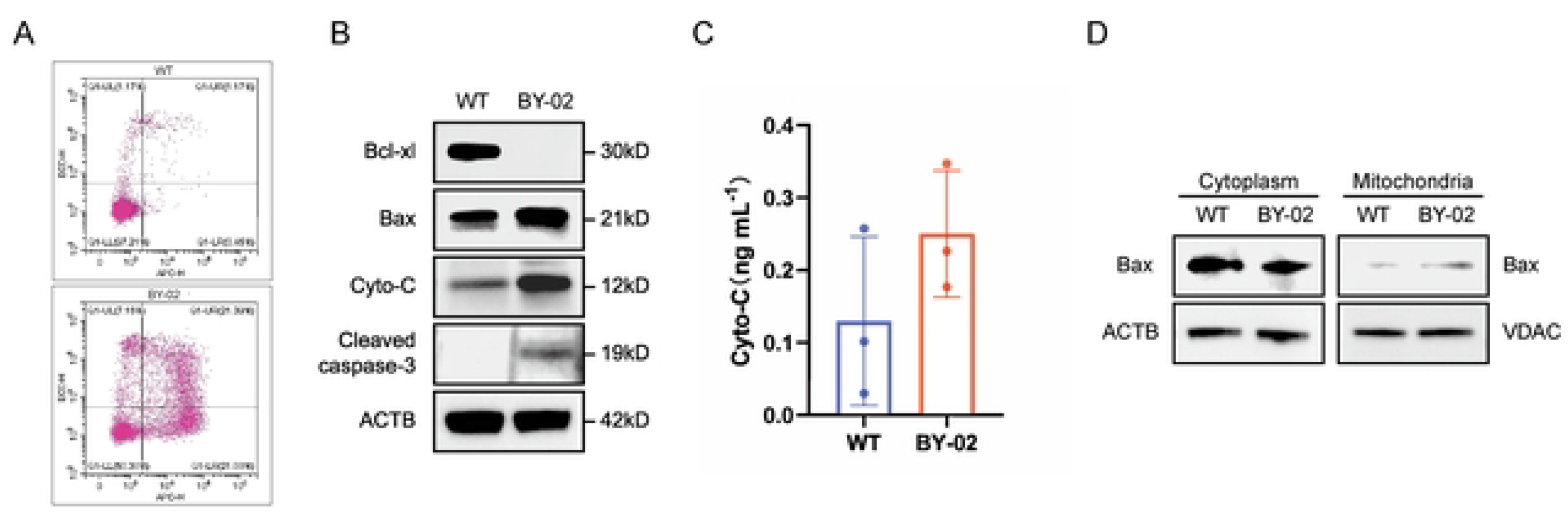
Effect of Bcl-xL on proteins involved in the apoptotic pathway of MDCK cells. (A) The level of apoptosis in BY-02 cells was analyzed by flow cytometry. The horizontal axis represents Annexin V-APC and the vertical axis represents PI. (B) Effect of Bcl-xL deficiency on apoptotic signaling pathways. ACTB protein was used as an internal control protein. (C) Cytochrome C content in the cell supernatant of WT and BY-02 cells. (D) The pro-apoptotic protein Bax content differed between WT and BY-02 cells in the cytoplasm and mitochondria. ACTB was used as a cytosolic reference protein and VDAC as a mitochondrial reference protein. Data are representative of at least three independent experiments.

### 3.5 Bioinformatics analysis of differentially expressed genes in BY02 cells

To further validate the tumor resistance of BY-02 cells in vitro and in vivo, the transcriptional level differences between BY-02 and WT cells were evaluated BY transcriptomics. DEGseq was used for differential analysis, A fold change >2 and a P value < 0.05 were used as the screening criteria for significant differential expression. In BY-02 cells, 687 genes were up-regulated and 260 genes were down-regulated (Fig. 5A). The differential gene expression clustering heatmap showed that the trend of gene transcription levels was the same in the three replicate samples of the same cell, and there were significant differences in gene transcription levels between BY-02 and WT cells (Fig. 5B). GO and KEGG analysis of differential genes showed that there were enrichment changes in cell migration and movement, angiogenesis and development, extracellular matrix components, signal transduction and cell structure (Fig. 5C). Enrichment analysis BY KEGG database showed that BY-02 cells mainly showed enrichment changes in related tumor signaling pathways and metabolism-related pathways (Fig. 5D). All these results point to the conclusion that Bcl-xL acts as a tumorigenic key gene in MDCK cells to promote their survival and development, and targeting Bcl-xL to inhibit its function is an effective solution to solve the tumorigenicity of MDCK cells.

**Fig. 5.**
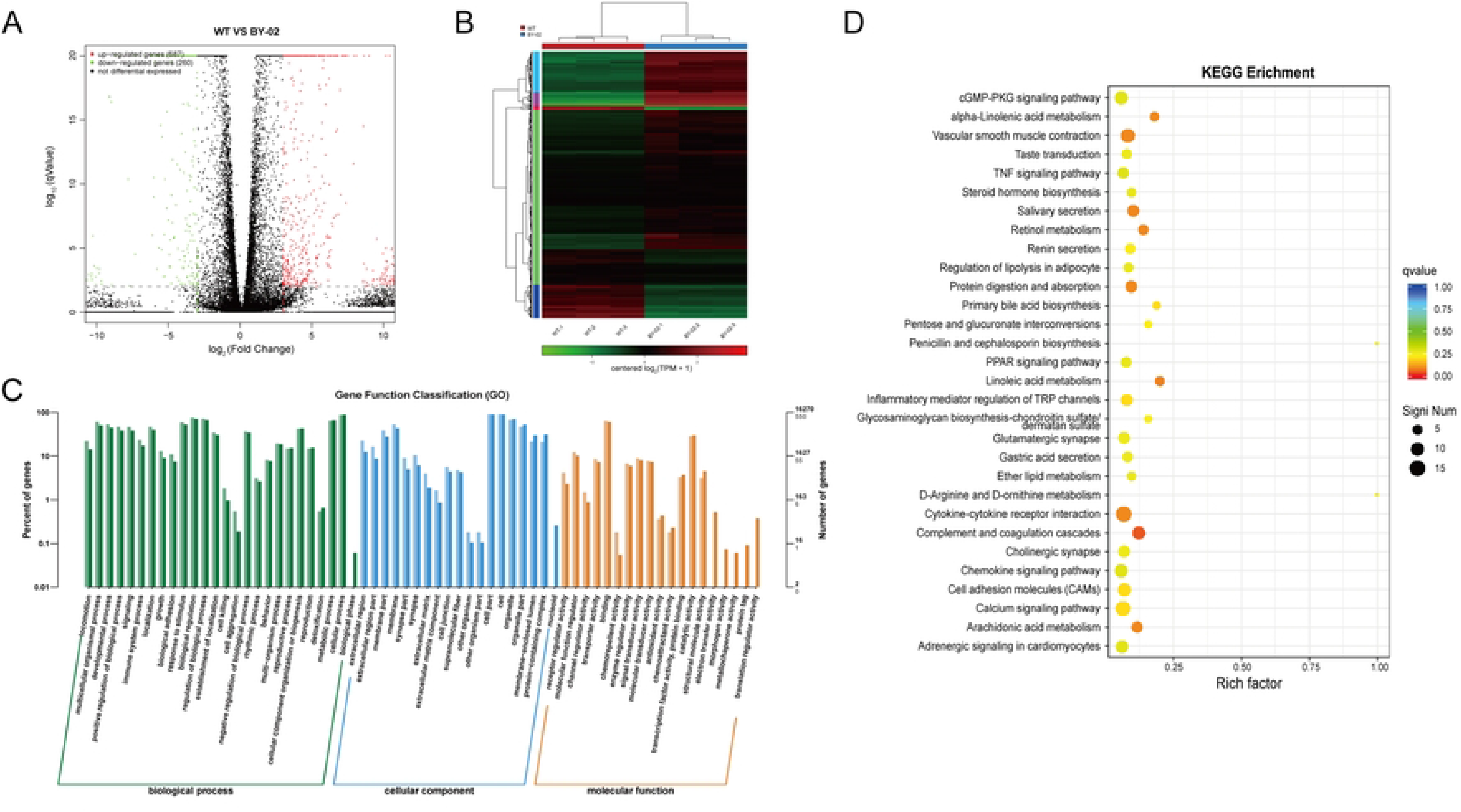
Analysis and comparison of transcriptome data between WT and BY-02 cells. (A) Volcano plots represent the significance and magnitude of changes in gene transcript levels between MDCK WT and BY-02 cells. Genes with significant differences are marked in red and green. Red represents upregulation and green represents downregulation. (B) Heat map of significantly altered proteins between WT and BY-02 cells. Color key indicates the relative abundance of proteins. (C) Differentially expressed genes between WT and BY-02 cells were subjected to GO enrichment analysis and classified into different categories based on biological process (BP), cellular component (CC), and molecular function (MF). (D) The effects of differential genes in the corresponding pathways in WT and BY-02 cells were analyzed.

## 4. Discussion

The rapid outbreak of SARS-CoV-2 underscores a significant challenge in vaccine development: the time and cost associated with production and preparation. Influenza, a disease that can spread between animals and humans, is no longer suitable for vaccine production using traditional egg-based methods. The World Health Organization supports the establishment of a new approach for influenza vaccine production, and the use of MDCK cell culture presents an attractive and alternative solution for influenza vaccine manufacturing. This method offers advantages, including the use of a cell bank capable of rapid exponential expansion and a stable virus passage without mutation. Influenza vaccines produced with MDCK cells are available in multiple countries. As a novel production cell line, the safety evaluation of MDCK cells holds significant importance, with a key focus on their potential for high tumorigenicity [26].To supply the demand, a MDCK cell strain with Bcl-xL deficiency, BY-02 was created in this work, which showed markable reduction of tumorigenicity in vivo.

Apoptosis plays a vital role in inhibiting tumorigenesis, and resistance to apoptosis is a defining characteristic of cancer. Mechanisms that enable cells to escape apoptosis are generally categorized as follows: (1) disruption of the balance between pro- and anti-apoptotic proteins, (2) reduced caspase activity, and (3) impaired death receptor signal transmission. The Bcl-2 protein family, comprising both pro- and anti-apoptotic proteins, holds great importance in regulating apoptosis, primarily at the mitochondrial level [27]. Dysregulated apoptosis in affected cells often results from an imbalance between anti- and pro-apoptotic proteins within the Bcl-2 family. This imbalance may involve the upregulation of at least one anti-apoptotic protein, the downregulation of at least one pro-apoptotic protein, or a combination of both [28]. Raffo et al. revealed that overexpression of Bcl-2 protects prostate cancer cells from apoptosis [29]. Fulda et al. found that Bcl-2 overexpression inhibits TRAIL-mediated apoptosis in neuroblastoma, breast cancer, and glioblastoma cells [30]. The dynamic balance between antiapoptotic and proapoptotic proteins is required to maintain homeostasis and cell survival [31]. After the cells are stimulated, Bax (the proapoptotic protein) is cloned into the mitochondrial outer membrane, which changes the mitochondrial permeability and releases cytochrome C to the cytoplasm [32] and activates Caspase 3, leading to the caspase cascade reaction and induction of apoptosis [33]. Overexpression of Bcl-xL can lead to the resistance of chondrosarcoma cells to conventional chemotherapy [34], which also occurs in colon cancer [35], liver cancer [36], and non-Hodgkin’s lymphoma [37]. Upregulation of Bcl-xL is linked to the multidrug-resistant nature of tumor cells, which hinders apoptosis [38].

Several studies have also shown that knocking down genes that encode anti-apoptotic proteins in the Bcl-2 family can increase apoptosis. For instance, pro-apoptotic and anti-proliferative effects of Bcl-2-specific siRNA have been observed in pancreatic cancer cells [39]. Thus, inhibiting anti-apoptotic proteins in the Bcl-2 family presents a feasible approach to preventing tumorigenesis. The strategy of this study was trying to promote apoptosis of MDCK cells to reduce tumorigenicity through knocking out anti-apoptotic gene *bcl-xl* using CRISPR-Cas9 system. MDCK cell lines with homozygous deletion of Bcl-xL were successfully screened using puromycin resistance tags. Compared with the WT MDCK cells, Bcl-xL knockout MDCK cells exhibited significantly decreased proliferation, colony formation, and migration ability. Flow cytometry analysis with Annexin V/PI staining suggested that Bcl-xL was crucial for MDCK cell apoptosis as deletion of Bcl-xL dramatically increased the level of apoptosis, which was 40% higher than that of WT MDCK cells. Further examination of the effect of Bcl-xL depletion on the key proteins related to the mitochondrial apoptotic pathway revealed that pro-apoptotic protein Bax was upregulated, and the downstream molecule cytochrome C was also detected in the cytoplasm and the supernatant. Simultaneously, the cleavage of apoptosis execution protein, Caspase 3, into the active fragment initiated apoptosis. Besides, Bax was translocated to the mitochondrial membrane in the absence of Bcl-xL, which was consistent with the previous reports that Bax aggregated on the mitochondrial membrane to produce cytochrome C by increasing membrane permeability, and induced apoptosis [40]. These results indicate that deletion of Bcl-xL be able to induce apoptosis in MDCK cells. Our subsequent studies showed that Bcl-xL deficiency remarkably suppressed carcinogenesis in MDCK cells with almost no tumorigenicity observed in nude mice. This indicated that a lack of Bcl-xL was able to inhibit MDCK cells from forming tumors.

Several years of research indicate that to overcome the limitations of chick embryo culture, the use of cell culture technology to isolate vaccine strains and produce vaccines is ideal. Currently, improving the virus titer and reducing the tumorigenicity of MDCK cells are the main directions to modify MDCK cells as a vaccine-producing strain. Thus, genetic engineering is expected to be a powerful tool for screening newly engineered cell strans. In short, this study provides a possible direction for the modification of MDCK cells with low tumorigenicity and lays a foundation for the construction of safer and more reliable cell lines.

## Institutional review board statement

All procedures were conducted in accordance with the “Guiding Principles in the Care and Use of Animals” (China) and were approved by the Ethical Review Committee of Experimental Animal Welfare, Wuhan Institute of Biological Products Co., LTD (WIBP-AII312023002).

## Data Availability Statement

Data will be made available on request.

## Funding and Acknowledgments

The present work was supported by the Ministry of Science and Technology of the People’s Republic of China (No. 2010AA022905) and Key Research and Development Program of Hubei Province “Novel influenza vaccine based on cell culture” (2020DCC002)

## Credit authorship contribution statement

Boran Li is a co-author of the paper. Changgui Li is the co-corresponding author of this paper. Jiayou Zhang, Zhegang Zhang, Daiguan Yu, and Changgui Li contributed to designing the research. Jiahao Zheng performed experiments. Jiahao Zheng, Lanxin Jia conducted animal experiments. Zheng Gong, Yang Le, Xuanxuan Nian, Xuedan Li, Bo Liu contributed to the result interpretation. All authors edited, revised, and approved the eventual manuscript.

